# Bridging the gap between in vitro and in silico single-molecule force spectroscopy

**DOI:** 10.1101/2022.07.14.500151

**Authors:** Diego E. B. Gomes, Marcelo C. R. Melo, Priscila S. F. C. Gomes, Rafael C. Bernardi

## Abstract

*Staphylococci* bacteria use an arsenal of virulence factors, mainly composed of proteins such as adhesins, to target and adhere to their host. Adhesins play critical roles during infection, mainly during the early steps of adhesion when cells are exposed to high mechanical stress. *S. epidermidis* SdrG:Fg*β* force resilience has been investigated using AFM-based single molecule force spectroscopy experiments paired with steered molecular dynamics (SMD) simulations. However, there is still a gap between both kinds of experiments at high force-loading rates. Here, we leveraged the high-speed of coarse-grained (CG) SMD simulations to bridge the gap between the data obtained *in vitro* and *in silico* with all-atom SMD. We used the DHS theory to connect the two types of SMD simulations and the predictions are consistent with theory and experimentation. We believe that, when associated with all-atom SMD, course-grained SMD can be a powerful ally to help explain and complement the results of single-molecule force spectroscopy experiments.

---

Mechanical forces play a fundamental role in biological systems. Experimental methods, such as Atomic Force Microscopy (AFM)-based single-molecule force spectroscopy (SMFS), allow for the direct measurement of mechanical properties of biomolecules. Meanwhile, *in silico* SMFS allows us to explore with atomistic detail mechanical properties of protein folds under mechanical stress.^1^ This methodology has been used by our group to investigate a myriad of systems, including cytoplasmic proteins,^2,3^ as well as extracellular proteins that are remarkably resistant to shear forces, including cellulosomal proteins,^4–7^ coronavirus proteins, ^8^ and bacterial adhesins.^9,10^ Beside those, we have also investigated the mechanical properties of the widely employed streptavidin-biotin complex. ^11,12^

Among the highly mechanostable protein complexes, adhesins are particularly interesting. *Staphylococci* adhesins play a crucial role during infection. They promote adhesion to the human extracellular matrix upon biofilm formation.^13,14^ Staph biofilms are associated with more than half of all nosocomial infections,^15^ with *Staphylococcus epidermidis* and *S. aureus* listed as the most common pathogens.^14,16^ The adhesion of these bacteria to the host exposes the cells to a high mechanical stress. ^17^

*Staphylococci* adhesins use a conserved “dock, lock, and latch” (DLL) mechanism—in which the host target, usually a peptide on the order of 15 residues, is first bound (dock), then buried (lock) between two immunoglobulin-like (Ig) fold domains N2 and N3,^18^ and finally a “latch” connects N3 back to N2, securing the complex (see **Figure 1**). Investigated earlier by our group, the interaction between the Serine-aspartate repeat protein G (SdrG) protein, from *S. epidermidis*, with the fibrinogen-*β* (Fg*β*) peptide was measured *in vitro* and *in silico* at the range of 2 nN, ^9^ being the highest stability among all non-covalent interactions, outperforming highly resilient protein complexes such as the cellulosome cohesion-dockerin type III interaction by a factor of four.^19^

**Figure 1:**
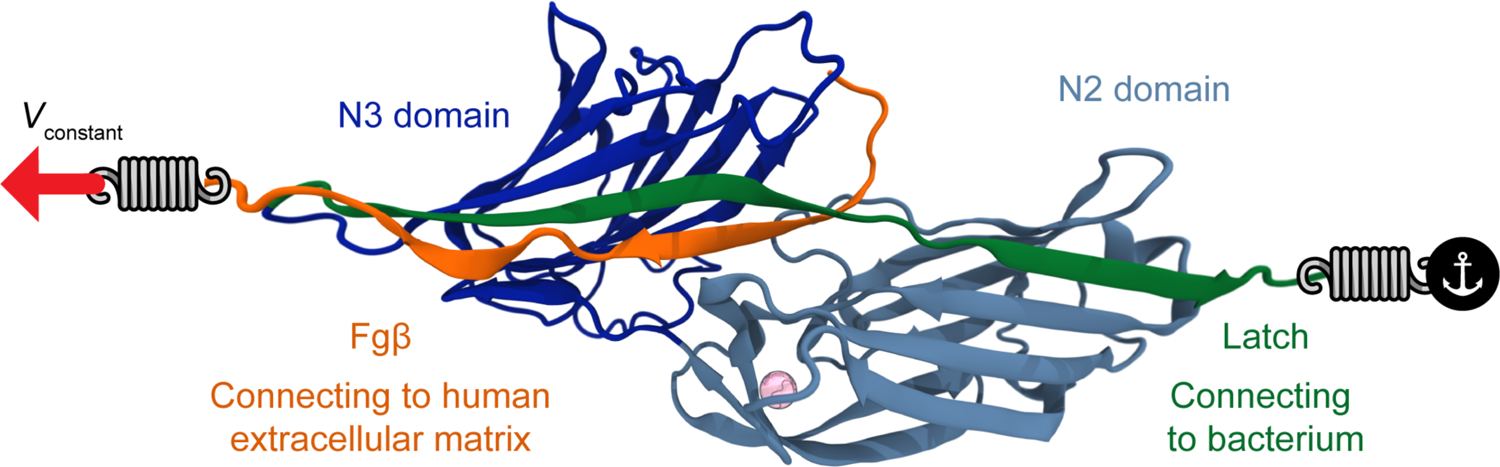
The SdrG:Fg*β* complex is shown in cartoon representations, the N2 domain is in light blue, N3 in dark blue, latch in green, and Fg*β* in orange. A structural Calcium ion is shown as a pink sphere. The figure indicates how the simulations were performed to resemble the mechanical strain imposed on the complex when in its biological context. The bacterial N2 domain was connected to a spring which in turn in anchored in a permanent position throughout the simulations. The peptide is also connected to a spring which is pulled away from the anchor point, at constant speed, pulling the complex apart. Protein rendering was built using VMD.^20^

Previously, Milles et al. used AFM-based SMFS experiments and all-atom steered molecular dynamics (aa-SMD) to explore the SdrG:Fg*β* complex under high-force regimes.^9^ Here we continued to explore the SdrG:Fg*β* complex mechanostability by leveraging the high-speed of coarse-grained MD (CG-MD) simulations to bridge the gap between the data obtained *in vitro* and *in silico*. The SdrG:Fg*β* complex presents an intricate behavior with load-dependent unbinding rates. In the SMFS experiments explored here, a molecular complex is pulled apart using increasingly faster pulling speeds, which in turn lead to increased loading rates. The higher loading rates accelerate dissociation, hindering thermal equilibration and allowing higher forces to accumulate before the rupture (**Figure 2**).

**Figure 2:**
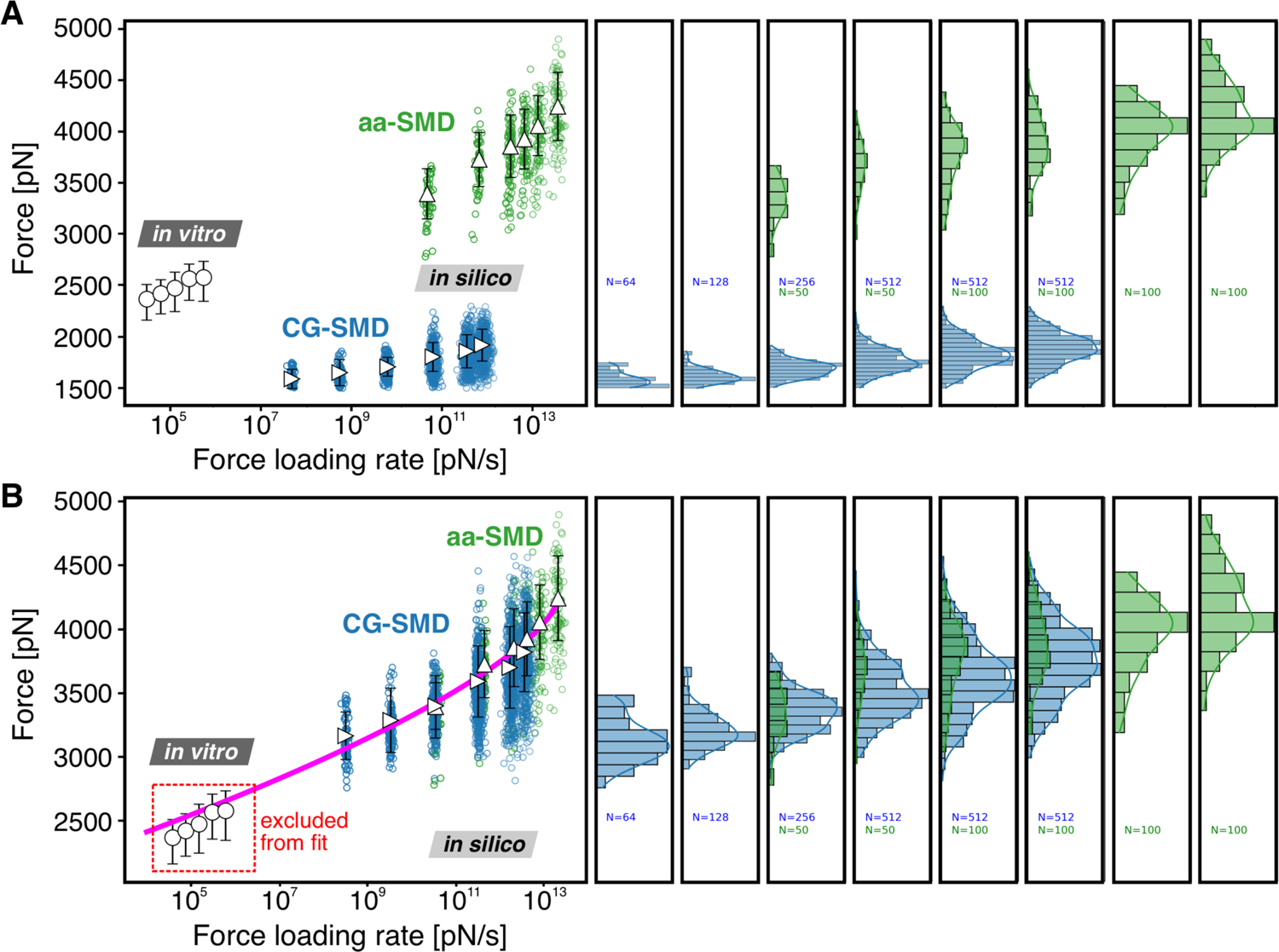
Dynamic Force spectrum for the SdrG:Fg*β* complex combining data from experimental SMFS, all-atom SMD and CG-SMD simulations. (A) CG-SMD simulations are presented before calibration with aa-SMD. (B) All-atom, and Coarse-grained steered molecular dynamics simulations (aa-SMD, and CG-SMD) were performed at different velocities: 2.5E-07 to 1.25E-02 nm/ps (green) and 2.5E-07 to 2.5E-03 (blue). A Dudko-Hummer-Szabo^21^ (DHS) fit was performed exclusively through the SMD data, including both the aa-SMD, and the CG-SMD. The DHS fits (Δ*x* = 0.111 nm, koff0 = 3.797E-22 *s*^-1^, Δ*G* = 75.8 kBT, magenta line) yield an adequate prediction of the experimental force results^9^ (shown as black open circles).

At extreme loading rates, a non-linear dependency emerges between the force required to split the molecular complex and the loading rate. To understand this complex behavior, a comprehensive theory for connecting dissociation rates from force spectroscopy experiments was presented by Dudko, Hummer, and Szabo (DHS).^21^ In essence, DHS use a diffusive barrier crossing interpretation to provide a probabilistic description for the most likely rupture force needed to dissociate a complex under a given loading rate. System-dependent parameters, such as the free energy barrier for dissociation (Δ*G*) for unbinding can be derived from experimental data using this formalism.

We used the DHS theory to connect two types of MD simulations, aa-SMD and CG-SMD, in a multi-scale approach, performed at multiple pulling speeds, and we show that the simulation predictions are consistent with theory and experimentation. The simulations were compared at different velocities and in several replicas (aa-SMD from 2.5E-07 to 1.25E-02 nm/ps, N = 500; CG-SMD from 2.5E-07 to 2.5E-03, N = 1984). The CG-SMD initially predicted much lower *most probable forces* and *most probable loading rates* compared to aa-SMD. This may be due to a combination of reasons, including harmonic bias intro-duced by the Gö-Model, the nature of the CG force field where multiple atoms or molecules contribute to a particle, or a longer 20 fs timestep. To account for this difference, we systematically re-scaled all four overlapping pulling rates (2.50e-05, 2.50e-04, 1.250e-03, and 2.50e-03 nm/ps) based on the fraction of *most probable loading rate* and *most probable force* of the lowest overlapping pulling rate (2.5e-05) between all-atom and coarse-grained simulations. Therefore, the coarse-grained loading rates and peak forces were multiplied by 6.92 and 1.99, respectively, before fitting data to the Dudko-Hummer-Szabo^21^ (DHS) model. The DHS fit (**Figure 2**) was performed exclusively through the SMD data, including both the aa-SMD, and the CG-SMD yielding a surprisingly adequate prediction of the experimental force results^9^ (Δ*x* = 0.111 nm, koff0 = 3.797E-22 *s*^-1^, Δ*G* = 75.8 kBT, magenta line).

In summary, we demonstrated the viability of a multi-scale *in silico* single-molecule force spectroscopy approach combining coarse-grained and all-atom representations, to predict the experimental behaviour of the ultra-stable SdrG:Fg*β* molecular complex, bridging the gap between simulations and experimental measurements, a milestone that lays the foundation for the application of multi-scale SMFS over a broader number of molecular complexes.

## Materials and methods

### All-atom simulations

The structure of the *S. epidermidis* adhesin SdrG binding to Fg*β* had been solved by means of X-ray crystallography at 1.86 Å resolution and was available at the protein data bank (PDB: 1R17).^18^ To increase the length of the peptide to mimic the one used experimentally, a longer peptide chain was randomly positioned following previously assigned protocols.^1,22^ Employing advanced run options of QwikMD,^23^ the structure was solvated and the net charge of the system was neutralized using sodium counter ions. In total, approximately 240,000 atoms were simulated in each simulation. The MD simulations in the present study were performed employing the GPU-accelerated NAMD molecular dynamics package.^24,25^ The CHARMM36 force field,^26^ along with the TIP3 water model^27^ was used to describe all systems. The simulations were performed assuming periodic boundary conditions in the NpT ensemble with temperature maintained at 300K using Langevin dynamics for temperature and pressure coupling, the latter kept at 1bar. A distance cut-off of 11.0 Å was applied to short-range non-bonded interactions, whereas long-range electrostatic interactions were treated using the particle-mesh Ewald (PME)^28^ method. The equations of motion were integrated using the r-RESPA multiple time step scheme^24^ to update the van der Waals interactions every step and electrostatic interactions every two steps. The time step of integration was chosen to be 2fs for all SMD simulations performed.

With structures properly equilibrated and checked, six different pulling speeds were used in constant-velocity SMD simulations (see **Table S1**), with slower velocity simulations having fewer replicas due to the 10-fold increase in computational cost. These simulations were previously published by our group,^29^ and are analyzed again here to calibrate a coarse-grained model.

### Coarse-grained simulations

The atomistic model of SdrG was modeled onto the Martini 3.0 Coarse-grained (CG) force field (v.3.0.b.3.2)^30^ using martinize2 v0.7.3.^31^ A set of native contacts, based on the rCSU+OV contact map protocol, was computed from the equilibrated all-atom structure using the server http://info.ifpan.edu.pl/~rcsu/rcsu/index.html32 and used to determine Gö-MARTINI interactions^33^ used to restraint the secondary and tertiary structures with the effective depth (*ϵ*) of Lennard-Jones potential set to 9.414 *kJ mol^-1^.* All CG simulations were performed using GROMACS version 2021.5.^34^

The SdrG+Peptide complex was centered in a rectangular box measuring with 10.0, 10.0, 25.0 nm to the X,Y, and Z directions. The anchor (SdrG C-terminal) and pulling (Peptide C-terminal) backbone (BB) atoms were used to align the protein to the Z axis. The box was then solvated with Martini3 water molecules. Systems were minimized for 10.000 steps with steepest descent, followed by a 10 ns equilibration at the NPT ensemble using the Berendsen thermostat at 298K, while pressure was kept at 1 bar with compressibility set to 3*e*^-4^ *bar*^-1^, using the Berendsen barostat. A time step of 10 fs to integrate the equations of motion. Pulling simulations were subsequently done at the NVT ensemble with a time step of 20 fs. The temperature was controlled using the v-rescale thermostat^35^ with a coupling time of 1 ps. For all coarse-grained molecular dynamics simulations, the cutoff distance for coulombic and Lennard-Jones interactions was set to to 1.1 nm,^36^ with the long-range coulombic interactions treated by a reaction field (RF)^37^ with *epsilon_r_* = 15. The Verlet neighbour search^38^ was used in combination with the neighbour list, updated every 20 steps. The LINCS^39^ algorithm was used to constrain the bonds and the leapfrog integration algorithm for the solution of the equations of motion. Several replicas of steered molecular dynamics simulations were performed at a range of speeds described in **Table 2**.

**Table 1:** SMD velocities and replicas. All velocities are given in nm/ps.

| Label | Velocity | Replicas |
| --- | --- | --- |
| V1 | 2.50e-05 | 50 |
| V2 | 2.50e-04 | 50 |
| V3 | 1.25e-03 | 100 |
| V4 | 2.50e-03 | 200 |
| V5 | 5.0e-03 | 100 |
| V6 | 1.25e-02 | 100 |

**Table 2:** SMD pulling rates and replicas for coarse-grained simulations. Pulling rates are given in nm/ps.

| Label | Velocity (nm/ps) | Replicas |
| --- | --- | --- |
| CG1 | 2.50e-07 | 64 |
| CG2 | 2.50e-06 | 128 |
| CG3 | 2.50e-05 | 256 |
| CG4 | 2.50e-04 | 512 |
| CG5 | 1.25e-03 | 512 |
| CG6 | 2.50e-03 | 512 |

## Acknowledgement

The authors thank Prof. Hermann Gaub and Dr. Lukas Milles for the helpful discussions. This work was supported by the National Science Foundation under Grant MCB-2143787 (CAREER: *In Silico* Single-Molecule Force Spectroscopy). The steered molecular dynamics simulations were performed at Blue Waters supercomputer as part of the Petascale Computational Resource (PRAC) grant “The Computational Microscope”, which is supported by the National Science Foundation (award number ACI-1440026 and ACI-1713784). Blue Waters sustained-petascale computing project was supported by the National Science Foundation (awards OCI-0725070 and ACI-1238993) and the state of Illinois. We thank Auburn University and the College of Sciences and Mathematics for the computational resources provided by Dr. Bernardi faculty startup funds.

